# Evolutionary dynamics of cytoplasmic segregation and fusion: mitochondrial mixing facilitated the evolution of sex at the origin of eukaryotes

**DOI:** 10.1101/034116

**Authors:** Arunas L. Radzvilavicius

## Abstract

Sexual reproduction is a trait shared by all complex life, but explaining its origin and evolution remains a major theoretical challenge. Virtually all theoretical work on the evolution of sex has focused on the benefits of reciprocal recombination among nuclear genes, paying little attention to the dynamics of mitochondrial genes. Here I develop a mathematical model to study the evolution of alleles inducing cell fusion in an ancestral population of clonal proto-eukaryotes. Mitochondrial mixing masks the detrimental effects of faulty organelles and drives the evolution of sexual cell fusion despite the declining long-term population fitness. Cell-fusion alleles fix under negative epistatic interactions between mitochondrial mutations and strong purifying selection, low mutation load and weak mitochondrial-nuclear associations. I argue that similar conditions could have been maintained throughout the eukaryogenesis, favoring the evolution of sexual cell fusion and meiotic recombination without compromising the stability of the emerging complex cell.

## 1. Introduction

Sexual reproduction with gamete fusion and reciprocal recombination is among the traits shared by virtually all eukaryotes (Ramesh et al., 2005; Goodenough and Heitman, 2014; Speijer et al., 2015). Current views on the evolutionary advantage of sex have it that recombination among nuclear genes exposes the hidden genetic variation in finite populations, breaks up unfavorable allele combinations under fluctuating selection, or rescues the genome from the mutational meltdown (Otto, 2009). These views, however, are based on the long-term effects of recombination, and do not directly explain when or how these traits first arose. It is becoming increasingly clear that sex first appeared as a part of the evolutionary transition from prokaryotes to eukaryotes, most likely after the endosymbiotic acquisition of mitochondria (Gross and Bhattacharya, 2010; Lane and Martin, 2010; Speijer et al., 2015). There is therefore far more than just recombination to the origin of sex, and a full account of the evolution of sexual reproduction has to account for the complex relationship between mitochondrial symbionts and the host.

Evolution of complex life can be conceptualized as a sequence of major evolutionary transitions (Buss, 1987; Maynard Smith and Száthmary, 1995). With each transition conflicts between the levels of individuality arise and have to be mediated for a stable higher-level unit to be established (Michod, 1997; Michod and Nedelcu, 2003). Conflict resolution often involves mechanisms reducing genetic variance within groups of lower-level units, eliminating the scope for defection and detrimental competition. In contrast, cell fusion allows for cytoplasmic mixing and horizontal spread of detrimental mutants, facilitating the evolutionary conflict and reducing the efficacy of purifying selection (Hastings, 1992; Bergstrom and Pritchard, 1998; Randerson and Hurst, 1999). The origin of cytoplasmic mixing at the early stages of eukaryogenesis therefore could have hindered the evolution of a stable higher-level — unit the eukaryotic cell (Radzvilavicius and Blackstone, 2015). While two mating types and uniparental inheritance (UPI) might eliminate these issues in modern eukaryotes (Birky, 1995; Hadjivasiliou et al., 2013; Sato and Sato, 2013; Greiner et al., 2015), the mechanism of asymmetric inheritance would not have been present during the early evolution of sex.

In their recent article, Havird et al. (2015) suggest a novel hypothesis for the evolution of eukaryotic sex, in which mitochondrial mutations play a central role. Owing to its high mutation rate, mitochondrial DNA (mtDNA) can quickly accumulate mutations compromising the function of the respiratory chain and diminishing cell’s viability, which prompts the evolution of compensatory nuclear modifications that could restore the cell’s fitness. Recombination among the nuclear genes would potentially increase the rate at which new compensatory combinations of nuclear alleles are introduced, rapidly improving the match between the two genomes. While important in many ways, the hypothesis does not account for one of the hallmark features of mitochondrial genetics—the cytoplasmic segregation (Rand, 2008, 2011)—which together with the purifying selection makes for the efficient elimination of mitochondrial mutations.

Another recent idea highlighting the role of mitochondria in the evolution of sex stems from the evolutionary history of endosymbionts and the biochemistry of cellular respiration (Blackstone and Green, 1999). Faced with stressful conditions constraining their growth and proliferation, proto-mitochondrial symbionts could have systematically manipulated the host cell phenotype using the by-products of oxidative phosphorylation. High emissions of reactive oxygen species (ROS), for example, could have served as a trigger for the host cell fusion and recombination, restoring favorable conditions for the endosymbiotic growth and proliferation. Similarly, for Speijer et al. (2015), mitochondrial acquisition gave rise to sex due to the ROS-induced genome damage and the need for frequent recombinational repair (see also Gross and Bhattacharya, 2010; Horandl and Hadacek, 2013).

While it is very likely that sex evolved in a cell that already possessed mitochondria (Lane and Martin, 2010; Lane, 2014; Speijer et al., 2015), the conditions favoring the emergence of cell fusion and cytoplasmic mixing under these circumstances have not received substantial attention. Multiple factors are likely to affect the evolutionary dynamics of mitochondrial mixing and segregation, including the intensity of purifying selection, mutation rate, epistatic interactions, intracellular competition and the properties of early cell cycles, but the relative importance of these effects in the early evolution of meiotic sex is not known. Here I introduce an infinite-population model to study cytoplasmic segregation at the origin of sexual cell fusion. I analyze the spread of proto-nuclear alleles inducing cell fusion with cytoplasmic mixing under the effect of purifying selection against mitochondrial mutations. The model suggests a set of conditions promoting the emergence of sex in the form of eukaryotic cell fusion, and strongly supports the view that mitochondria could have represented one of the driving forces behind the origin of sexual life cycles.

## 2. Mathematical model for cytoplasmic segregation and the evolution of sexual mixing

### 2.1 Neutral segregational drift with clonal host reproduction

The eukaryotic cell can be modelled as a collective of mitochondria within a cytosol which also contains the host genome (I assume here that similar conditions also applied early in eukaryogenesis, i.e. after the acquisition of mitochondria). Consider an infinite population of cells, containing M mitochondria each and reproducing clonally. Mitochondria are found in one of two possible states, wild-type or mutant. The clonal reproduction is modelled by first duplicating the mitochondrial population of the cell and then randomly partitioning organelles to the two daughter cells through random sampling without replacement A cell containing *M* mitochondrial mutants will give birth to a daughter with *q* mutations with probability

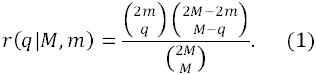

The frequency distribution for the number of mutants per cell *p(x)* after one round of clonal reproduction will therefore change to 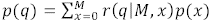. The process of cytoplasmic segregation modeled in this manner represents a type of neutral genetic drift, conceptually similar to the Wright-Fisher process (Ewens, 2004), but assuming a finite group size at all stages of the life cycle.

For the initial frequency of mitochondrial mutations within the cell *f_0_*=*m_0_/M*, variance after *n* clonal divisions is (see Appendix for the details)

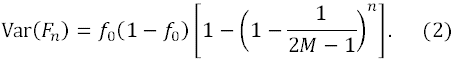

Segregational drift therefore increases variance between host cells, which after a large number of clonal reproduction cycles converges towards *f*_0_(1 − *f*_0_), where all mutants have either reached fixation within the cell or were replaced by the wild type mitochondria. It follows then that the probability to reach fixation is equal to the initial mutant frequency *f*_0_.

It can be similarly shown (Appendix) that neutral segregation increases the homogeneity within the cell. The probability for two segregating units within the cell to be identical by descent can be expressed as

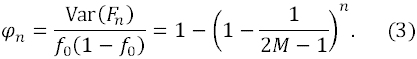

After a large number of clonal divisions, the identity-by-descent probability approaches one, *φ* → 1, at which point the mitochondrial populations are fully clonal and no further change is possible.

### 2.2. Evolution of cytoplasmic mixing

Now consider a full population life cycle with mitochondrial mutation, selection and reproduction (Fig. 1). We assume an ancestral state without sex or cell-cell fusion, where haploid hosts reproduce clonally as described above. The mode of reproduction is controlled by a single locus in the host’s haploid genome, *h/H*. Only mutants carrying a copy of the allele *H* can initiate a temporary cell fusion with a randomly chosen partner (3a-3c in Fig. 1) before proceeding to the standard clonal reproduction.

**Figure 1.**
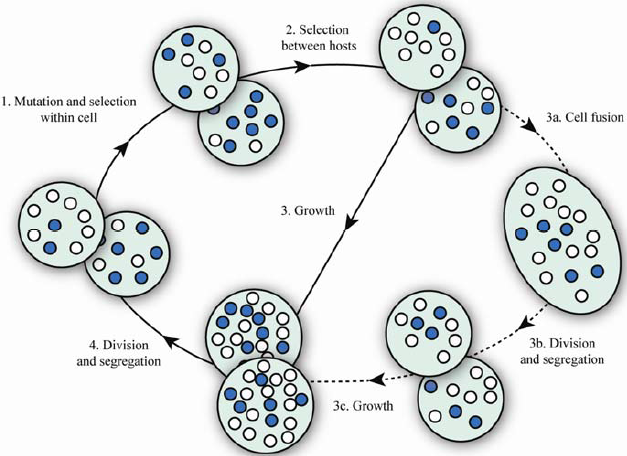
Population life cycle with asexual and sexual modes of reproduction. Shaded circles represent deleterious mitochondrial mutants within the host cell, cooperative organelles are left blank. Steps 1–4 (solid arrows) represent the life cycle of clonally reproducing individuals. Steps 3a–3c (dashed arrows) occur only if one of the cells meeting at random is a carrier of the cell fusion allele *H*.

The population state at generation *t* is represented by a (*M* + 1) ⨯ 2 matrix **P**^*(t)*^ with the matrix element 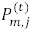 depicting the frequency of cells with *m* mitochondrial mutants and the nuclear state *j* (*j* = 0,1). The column vector 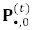 therefore corresponds to the wild type population with allele *h* and column 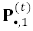 contains entries pertaining to the cells with the nuclear allele *H*.

#### 2.2.1 Mutation

The genotype-changing events in the population life cycle can be represented as matrix operations changing the population state **P**(^t^). The population state after the mutation is therefore given by **P**^(*t*,1)^ = **UP**^(*t*)^, where **U** is (*M* + 1) ⨯ (*M* + 1) transition matrix, with the element *U_ij_* defined as a probability that a cell with *j* mutant mitochondria will contain *i* mutants after the transition. Mitochondrial mutation at the rate *μ* is modeled as a binomial event, giving the transition probabilities

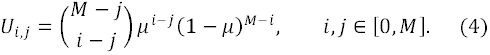

#### 2.2.2. Selection on the lower level

In the case where mutant mitochondria have a competitive advantage within the cell (“selfish” mutants), the mutation step is followed by selection on the lower level. Selection among mitochondria of the same cell is modeled as a random sampling with replacement, with the probability to select a selfish mutant proportional to its replicative advantage 1 +*k*. The population state after selection on the lower level is therefore **P**(*t*,2) = **WP**(*t*,1), where the binomial transition probabilities of matrix **W** are

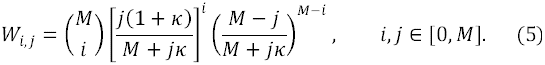

#### 2.2.3. SElection between eukaryotic hosts

Selection on the higher level changes the relative genotype frequencies according to the host cell fitness. In matrix notation the population state after selection is

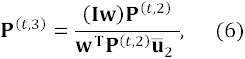
where **I** is the identity matrix, 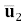 is a column vector of ones (1,1)^T^, and **W** is a column vector with the m-th element *w_m_* = ω*(m)* corresponding to the fitness of a cell containing *m* mutants. Following the models by Hadjivasiliou et al. (2013) and Kuijper et al. (2015), I assume that the relative fitness of the cell depends on the number of mitochondrial mutants *m* and can be expressed as *ω_(m)_* = 1 − *s(m/M)*^ξ^. Parameter *s* here represents the strength of selection and ξ determines the strength of epistatic interactions between mitochondrial mutations. Empirical studies suggest that in modern eukaryotes ξ > 1, leading to the so-called phenotypic threshold effects (Rossignol et al., 2003). The relative effect of each new mutation therefore increases with the overall mutation load.

#### 2.2.4. Reproduction and mitochondrial segregation

Cells carrying the nuclear allele *H* are capable of cytoplasmic fusion with the other *H-* type individuals as well as randomly chosen wild-type hosts *h*. Wild-type individuals do not initiate cell fusion, and mix their cytoplasmic contents only if randomly chosen by an individual carrying the allele *H*. The process of cell fusion in our model is represented by the convolution of corresponding genotype-frequency vectors, forming a temporary subpopulation of diploid zygotes each containing *2M* mitochondria. Fusion is immediately followed by cell division with random partitioning of mitochondria between the two daughter cells. The population state after the sexual stage of the life cycle can then be expressed as

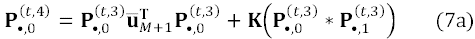

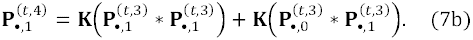

Asterisks here denote vector convolution, and 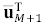 is a row vector of *M* + 1 ones, so that 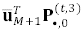 is the total frequency of the allele *h*. **K** is the transition matrix for the reductive cell division without the prior replication of mitochondria, implemented as selection without replacement with transition probabilities (Eq. 1)*K_i,j_* = *r(i∣M,j/2)*, where *i* ∈ [0,*M*],*j* ∈ [0,*2M*].

The life cycle ends with the standard clonal replication, first duplicating the mitochondrial population within each cell and then partitioning the organelles between the two daughter cells. This gives the updated population state at the start of the next generation **P**^(*t*+1)^ = **SP**(*t*,4), with transition probabilities (Eq. 1) *S_i,j_* = *r(i∣M,j)*, where *i,j* ∈ [0, *M\*.

The model is initialized in a random mitochondrial state **P**^0^ so that the whole population initially consists only of the wild-type individuals, i.e.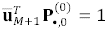 and 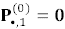. After the equilibrium is reached at time *t_E_*, the allele *H* is inserted at a small frequency χ = 0.001, so that 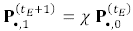 and 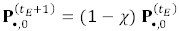. In the following **I** present the results based on the numerical solution of the above system of equations for both equilibrium and transient states, obtained for multiple values of *μ,k,s,M* and *ξ*.

## 3. Mitochondrial variation and the fitness cost of cytoplasmic mixing

### 3.1. Mitochondrial Variation at the Segregation-Fusion Equilibrium

Let us first look into the effect of recurrent cell fusion on the mitochondrial variance between cells generated by cytoplasmic segregation. Starting with a fixed number of mutants per cell *m_0_* **I** allow for *η* clonal generations without cell fusion, followed by a single round of sexual reproduction. No mutation or selection occurs at this stage.

The results show that the effect of cytoplasmic mixing opposes the constant increase of mitochondrial variance between cells generated by segregational drift (Eq. 2), establishing an intermediate equilibrium (Fig. 2). While segregational drift alone results in diverging cell lineages with clonal intracellular populations of mitochondria, increasing frequency of sexual cell fusion relative to the number of clonal generations *η* reduces the mitochondrial variation between cells, at the same time reducing homogeneity within the cell. Given the importance of heritable variance for the efficacy of selection on the higher level, frequent cell fusion could result in diminished population fitness, which I investigate further.

**Figure 2.**
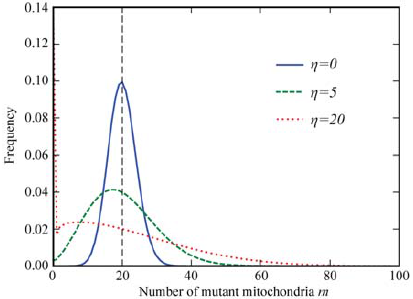
Cytoplasmic fusion opposes the effect of mitochondrial segregation of neutral mutations. Variance in the number of mutant mitochondria per cell increases in multiple rounds of clonal reproduction, but is reduced by cell fusion, resulting in an equilibrium when the two modes of reproduction alternate in time. *η* is the number of consecutive clonal generations without cytoplasmic mixing. The initial number of mutants per cell is *m_0_ = 20* (vertical line).

### 3.2 Mitochondrial mutation pressure

Here I return to the full population life cycle and analyze the effects of cellular fusion on the long-term population fitness. Mitochondrial mutants arise at a constant rate *μ* but do not have an intra-cellular replication advantage over the cooperative organelles, i.e. *k* = 0. Cell fusion rate is controlled by keeping the frequency of nuclear allele *H* at a constant level *p_H_*, while allowing the mitochondrial population to evolve freely. Given that a cell carrying the *H* allele fuses with a randomly selected partner, the overall rate of sexual reproduction can be expressed as 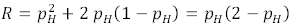.

Owing to the effect of reduced variance in the number of mitochondrial mutants between cells, the long-term population fitness is reduced by increasing frequency of *H* (Fig. 3). In agreement with previous studies (Hadjivasiliou et al. 2013, Radzvilavicius et al., 2015), the detrimental effect of lower mitochondrial variance is more prominent with higher numbers of mitochondria per cell. Larger populations of segregating lower-level units dampen the effect of segregational drift (Eq. 2), reducing the efficacy of selection on the higher level.

**Figure 3.**
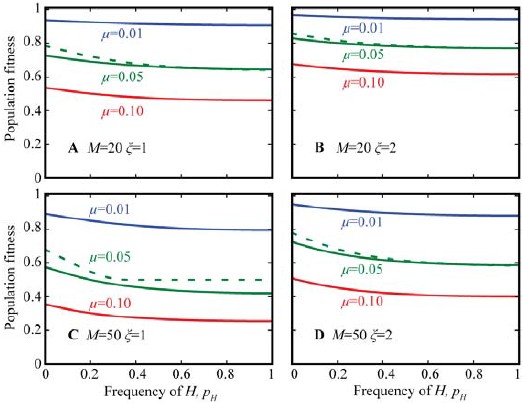
Frequent cytoplasmic mixing reduces the mean population fitness under mitochondrial mutation pressure. *H* is the cell fusion allele. *M* is the total number of mitochondria per cell,μ is the mitochondrial mutation rate and ξ is the strength of epistatic interactions. Selection strength is set to *s* = 1, except for μ = 0.05, where dotted lines show the effect of weaker selection with *s* = 0.5.

### 3.3. Fast replicating “selfish” mutants

Fitness costs of cytoplasmic mixing can be exacerbated in the presence of so called “selfish” mitochondria—organelles that have gained mutations leading to the faster reproduction rate but at the same time reducing their ability to participate in cooperative interactions. Selection on the lower level therefore increases the frequency of non-cooperative mitochondria, but cells with a significant proportion of selfish lower-level units suffer a fitness cost and replicate slower. This is an example of the evolutionary conflict between levels of selection, endangering the stability of the higher-level unit—the nascent eukaryotic cell (Schable and Wise, 1988; Taylor et al., 2002; Clark et al., 2012; Bastiaans et al., 2014). This time, I assume that mutants arise at a low rate (*μ* = 0.0001) and proliferate mostly due to their ability to outcompete the cooperative mitochondria within the same cell.

The results confirm that clonal reproduction (low*p_H_*) coupled with purifying selection on the higher level suppresses selfish mitochondrial competition within the cell and maintains high population fitness (Fig. 4). Unable to spread horizontally in the absence of cell fusion, selfish mutants affect only those cell lineages in which rare mutations occur, allowing selection on the higher level to rapidly eliminate the affected individuals. There is, however, a critical frequency *p_H_* at which the non-cooperative mitochondria start proliferating faster (Fig. 4A-F). Frequent cytoplasmic mixing reduces the mitochondrial variance between cells and therefore lowers the efficacy at which selection can eliminate the affected cells. The conditions are more permissive for selfish proliferation in populations with high numbers of mitochondria per cell (Fig. 4A-D), weak selection (Fig. 4E-F) and strong epistasis *ξ* (Fig. 4E-F). With increasing *ξ* the fitness function becomes increasingly flat at low *m*, allowing the non-cooperative mitochondria to reach high per-cell frequencies before they are eliminated by selection.

**Figure 4.**
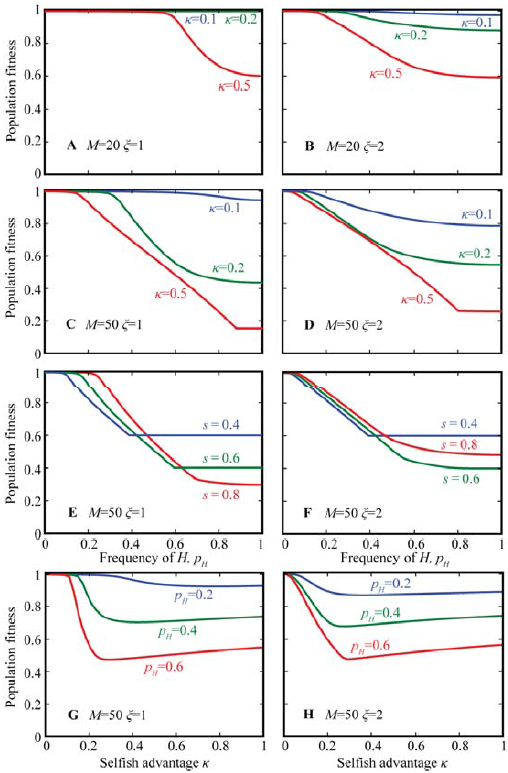
Fitness costs of cytoplasmic mixing can be exacerbated in the presence of selfish mitochondrial mutants. Cooperative interactions within groups of mitochondria break down easier in larger groups (A and C vs. B and D) with strong epistasis (A, C, E and G vs. B, D, F and H) and weak selection on the higher levels (E–F). Excessive competition among mitochondria can become costly to selfish organelles, increasing the mean population fitness, if fast replicating deleterious mitochondria overtake the cell before being able to spread (G–H). Mutation rate is set to μ = 0.0001, selections strength *s* = 1 unless indicated otherwise. Selfish advantage is set to *k* = 0.2 in E and F. *H* is the cell fusion allele, ξ is the strength of epistatic interactions.

Further increase in *p_H_* pushes the equilibrium towards the lower population fitness by increasing both the number of mutants per cell and the amount of affected lineages. Eventually all cell lineages contain some non-cooperative mitochondria at which point the population fitness becomes nearly independent of the frequency of *H* (Fig. 4A-F).

Interestingly, the equilibrium frequency of selfish mitochondrial mutants is not always a monotonic function of their relative replicative advantage *K* (Fig. 4G-H). There is a critical value of *K* correpsonding to the highest mutant load and the minimal population fitness. With the replicative advantage lower than the critical value of *k*, the mutant spread through the population is limited by their replication rate; for higher *K*, selfish mutants overtake their host cells too rapidly, allowing the purifying selection to efficiently suppress their further spread.

## 4. Invasion of *H* mutants

In this section I consider an evolutionary scenario where alleles *H* are introduced into a population at a low frequency and evolve freely. The allele *H* changes the mode of reproduction by inducing temporary cell fusion with a randomly selected partner, mixing the mitochondrial populations of the two cells (Fig. 1).

### 4.1. Epistasis between mitochondrial mutations

The results show that the cell-fusion allele *H* is able to invade, spread to an equilibrium frequency of *p_H_* < 1 or reach fixation (*p_H_* = 1) (Fig. 5A). The invasion occurs in spite of the curtailed long-term population fitness due to the lower variance in the number of mitochondrial mutants among invaders and weaker selection (Fig. 5B). The necessary condition for successful invasion is *ξ* > 1, i.e. the negative epistasis between deleterious mitochondrial mutations. The detrimental effect of every new mutation has to increase with the total mutational load, as indeed is the case in modern eukaryotes.

**Figure 5.**
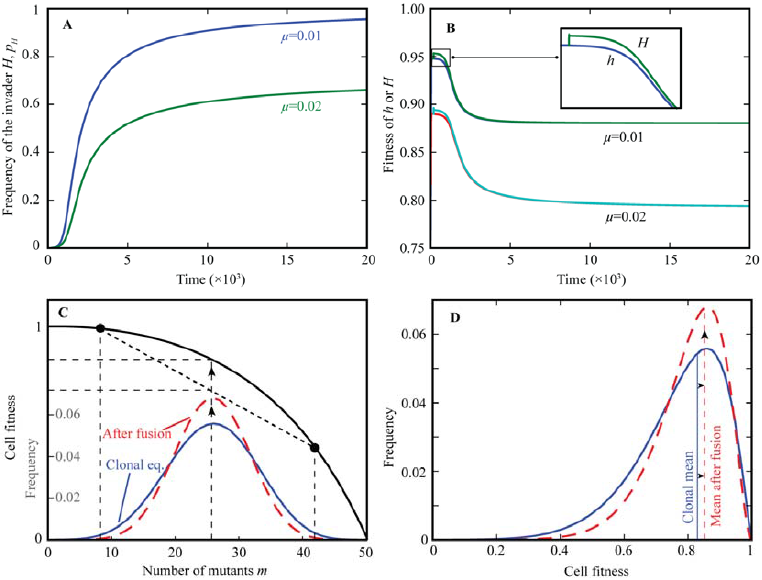
Invasion of the cell fusion allele *H* into an ancestrally clonal population. A nuclear allele inducing cytoplasmic fusion invades and can reach fixation (A), but its spread reduces the long-term population fitness (B). A singular cell fusion-division event increases the frequency of intermediate cytotypes, and reduces the frequency of cells with extreme mutant numbers (C). The loss of mitochondrial variance reduces the efficacy of selection and is detrimental in a long term. However, the reduced frequency of extreme cytotypes can be beneficial in a short term (D), if the intermediate cytoplasmic states have a higher fitness than expected from the additive interactions, i.e. with negative epistasis (*ξ* > 1) (C). The fitness advantage can be maintained if the mito-nuclear linkage is weak, e.g. if half of the mitochondria are inherited from a randomly selected partner which would otherwise reproduce clonally. *μ* = 0.04,*ξ* = 3 in C and D, *ξ* = 2 in A and B. The number of mitochondria per cell is *M* = 50,*k* = 0.

Cytoplasmic mixing increases the frequency of intermediate cytoplasmic states, reducing the frequency of cells with extreme mutant numbers (both high and low, Fig. 5C). This reduced variance has a long-term fitness disadvantage due to the weakened response to selection (Fig. 5B). However, with negative epistasis (*ξ* > 1), the intermediate cytotypes have a higher fitness than expected from the linear combination of the extremes, which gives the invading allele *H* a short-term advantage (Fig. 5D). Invaders choose their mating partners randomly and therefore the mitochondrial-nuclear associations remain weak. This allows the allele *H* to acquire a long-lasting advantage over the clonally reproducing subpopulation *h* (Fig. 5B), even though its spread inevitably curtails the population fitness in the long term. The advantage is lost with positive epistasis (*ξ* < 1), in which case both short-and long-term effects of reduced mitochondrial variance become detrimental.

### 4.2. Further conditions favoring the spread of H

Under the pressure of deleterious mitochondrial mutations, the allele *H* spreads to high frequencies and fixes more readily with low mutation rates (Fig. 6A-B). Cytoplasmic fusion has a stronger evolutionary advantage with small mitochondrial population sizes, as segregational drift is more efficient in generating mitochondrial variance with small *M* (Eq. 2). A similar trend is observed with selfish mitochondria having a replicative advantage over their wild-type counterparts, where fast replication of mutants, i.e. large *K*, diminishes the evolutionary advantage of the cell-fusion allele *H* (Fig. 6C-D). This time, there is a critical value of *K* corresponding to a distinctive drop of equilibrium allele frequency *p_H_* to zero. This fast transition occurs once the replicative advantage of selfish organelles becomes large enough to rapidly reduce the fitness of fusing hosts, whereas the high-fitness asexual lineages remain resistant to their spread. As the allele *H* spreads due to its short-term fitness advantage, its fixation is also facilitated by strong purifying selection on the higher level *s* (Fig. 6E-F).

**Figure 6.**
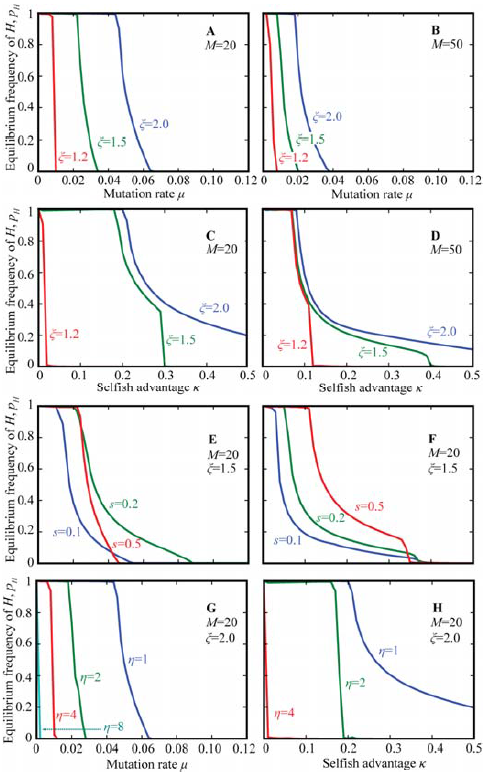
Conditions favoring the evolution of sexual cell fusion. Reduced mutation rates, small mitochondrial populations and negative epistatic interactions all promote the evolution of sexual cell fusion under mitochondrial mutation pressure (A–B). In the presence of selfish mitochondrial mutants, competition on the lower level has to be suppressed before cell fusion can be established (C–D). Cellular fusion evolves easier under strong purifying selection on the higher level (conditions favoring the fixation of *H* are less strict under large *s*, E–F). With alternating clonal and sexual life cycle stages, the number of consecutive clonal divisions *η* must remain low (G–H). Mutation rate in C, D, F and H is set to μ = 0.0001,*s* is the strength of selection, *H* is the allele inducing cell fusion with a randomly selected partner.

### 4.3. Alternating life cycles and mitochondrial-nuclear associations

Consider now the case where mutants carrying the allele *H* are capable of inducing the cytoplasmic fusion only every *η* generations. It is indeed often the case in protists that individuals engage in sexual reproduction only occasionally, e.g. under stressful environmental conditions or starvation (Dacks and Roger, 1999; Goodenough et al., 2007). Since the clonal stage of the life cycle would result in higher variance due to the segregational drift, *η* is likely to affect the evolutionary success of the cell fusion allele *H*. Indeed, the model shows that increasing number of consecutive clonal divisions in-between the sexual fusion events has a very strong effect *opposing* the spread of the fusion allele *H* (Fig. 6G-H). As few as 4–8 clonal cell divisions could be enough to prevent the invasion of the cell fusion allele *H* under all reasonable conditions investigated.

The principal reason allowing the allele *H* to invade lies in the way reduced mitochondrial variance affects the mean fitness after every cellular fusion event Reduced variance in the number of mutants gives a fitness advantage due to negative epistasis, but only if the association between the nuclear allele and the mitochondrial population of the same cell is temporary. This is easiest achieved through frequent fusions with randomly chosen partners that might otherwise reproduce clonally. With *η* = 1 mitonuclear associations are weakest, but become stronger when the same mitochondrial population persists within the lineage for several generations, i.e. *η* > 1. Detrimental long-term effects of reduced mitochondrial variation between the higher-level units become increasingly important as *η* grows. Fixation of the cellular fusion allele *H* therefore requires frequent cytoplasmic mixing, maintaining weak mitochondrial-nuclear associations.

## 5. Conclusions and discussion

It is very likely, that sex appeared in a cell that already possessed mitochondria, but not mating types or mechanisms constraining the cytoplasmic inheritance. The model presented here shows that sexual cell fusion can evolve simply to promote mitochondrial mixing, which temporarily covers up the detrimental effects of faulty or selfish endosymbionts or organelles.

Low mitochondrial mutation rates allow cell fusion alleles to fix more readily (Fig. 6A–B). Admittedly, substantial variation in mitochondrial mutation rates among the extant eukaryotes makes the inference of the ancestral pace of mutation accumulation rather complicated. On the one hand, we know that evolution rates (a proxy for the true mutation rate) in intracellular symbiont genomes are typically elevated (Itoh et al., 2002; Marais et al., 2008). Mitochondrial evolution rates in higher animals, fungi and some plants can also be substantially higher than in their nuclear genomes (Lynch et al., 2006, 2008; Sloan et al., 2009). On the other hand, the mutation rate appears to be extremely low in most plants, early branching metazoans (Palmer and Herbon, 1988; Shearer et al., 2002; Huang et al., 2008) and many unicellular eukaryotes (Burger et al., 1995, 2013; Smith and Keeling, 2015). It is therefore not impossible that the initially high evolution rate at the beginning of the endosymbiotic association slowed down as the evolutionary transition progressed, facilitating the evolution of sex. High mutation rates in some present-day eukaryotes are then secondarily derived, perhaps owing to their high metabolic rates and active lifestyles.

While the initial symbiotic association remains shrouded in mystery (Martin and Muller, 1998; Embley and Martin, 2006; Martin et al., 2015), with mitochondrial endosymbiosis entering an obligatory phase, selection against mitochondrial mutations, e.g. the ones affecting the respiratory function of the cell, likely increased in strength (higher *s*) providing conditions more permissive for the evolution of cytoplasmic mixing (Fig. 6E-F). Indeed, empirical data reveal substantial purifying selection acting on mitochondrial populations in modern animals (Elson et al., 2004; Stewart et al., 2008; Castellana etal., 2011, Cooper et al., 2015). Similarly, as the evolutionary transition progressed, selfish mitochondrial competition also had to become suppressed (reduced *K*), through the proto-eukaryotic mechanisms of conflict mediation, e.g. honest signaling, membrane uncoupling and reduced mitochondrial genomes (Radzvilavicius and Blackstone, 2015), thus promoting the evolution of meiotic sex (Fig. 6C–D).

Extrapolating from fitness interactions in modern eukaryotes (Rossignol et al., 2003), it is not unreasonable to assume negative epistatic interactions (*ξ* > 1) between detrimental mitochondrial mutations throughout the eukaryogenesis, favoring the invasion of cell-fusion alleles (Fig. 6A). Indeed, with multiple mitochondria per cell and several copies of mtDNA per organelle, a critical amount of deleterious mutations has to accumulate before cellular respiration is significantly impaired (Mazat et al., 2001; Rossignol et al., 2003). Mitochondrial endosymbiosis therefore created a unique genetic system in which strong synergistic interactions between deleterious mutations favor the evolution of sexual cell fusion. This is in stark contrast to the deleterious mutations in the nucleus (or bacterial genomes), where negative epistatic interactions are relatively uncommon (Kouyos et al., 2007).

Modern eukaryotes are capable of reproducing clonally in numerous consecutive generations, punctuated by occasional sex (Dacks and Roger, 1999; Goodenough et al., 2007). The model analyzed here predicts, however, that when regulated cell fusion first arose, it had to be frequent—and clonal reproduction rare—in order to maintain weak mitochondrial-nuclear associations responsible for the evolutionary advantage of cytoplasmic mixing, i.e. low values of *η* (Fig. 6G-H). Frequent cell fusion events might have been vital early in eukaryogenesis, since without the precisely coordinated chromosome and cell division machinery, consecutive reproduction cycles without cell fusion could have produced nonfunctional chromosome numbers or gene combinations, rendering the emerging eukaryotic cell unviable. Similarly, frequent cell fusions would have masked mutations and stabilized the emerging eukaryotic genome in the presence of intron bombardment and endosymbiotic gene transfer (Lane, 2011).

One way to interpret the main result of this work is that the initial selective pressure driving the evolution of cell-cell fusion could have been mitochondrial, in which case the routine recombination among nuclear genes came as a fortunate side effect, maintaining the evolutionary advantages of sex past the evolution of the uniparental inheritance and until present day. Indeed, the molecular machinery for the meiotic, reciprocal recombination had to evolve in the routine presence of cell fusion events, and the barrier separating the eukaryotic sex from prokaryotic recombination might have never been crossed without mitochondria. It is more likely, however, that multiple mechanisms promoting cell fusion acted at the same time, with mitochondrial selection pressure contributing to the ease at which sexual reproduction with cytoplasmic fusion and reciprocal recombination came into the widespread existence.

## Acknowledgements

Andrew Pomiankowski, Nick Lane and Neil W. Blackstone are thanked for their contributions to this work. The author is supported by a UCL CoMPLEX PhD award from the Engineering and Physical Sciences Research Council (EP/F500351/1).

## Appendix Variance and identity-by-descent relations

Let*X_n_* be a random variable denoting the number of mutants within a cell after *n* rounds of clonal cell division, sampling without replacement *M* mitochondria from the doubled population of *2M*. The population mean is then simply equal the initial number of mutants within the cell, E(*X_n_*) = *x*_0_. Variance in the number of mutants can be expressed as

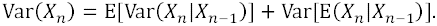

Given that the variance of the hypergeometric probability distribution used in sampling without replacement is

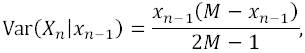

we can further write

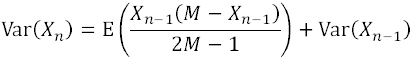

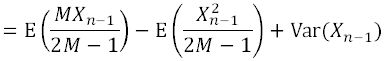

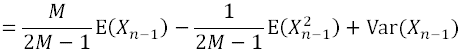

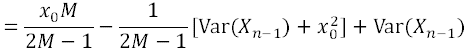

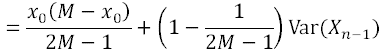

Here *x_0_* is the initial number of mutants within a cell. With the boundary condition Var(*X*_0_) = 0 the solution is

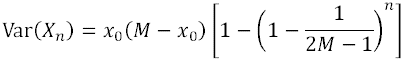

Variance in the mutant frequency 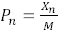 is then

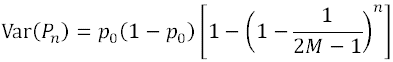

The genetic diversity (or lack of it) within a cell can be expressed as a probability that two randomly selected mitochondria within the cell will be identical by descent,*f_n_*. In our random segregation model two lower-level units are considered identical if they are either descendants of the same parent, or different parents that are identical by descent themselves due to associations in previous generations. After *n* generations we can then write

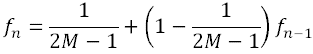

The above recursion is straightforward to solve for the parameter of non-identity *h_n_* = 1 − *f_n_*. As initially all mitochondria within the cell are assumed to be unrelated, the boundary condition is *h_0_* = 1. It is then easy to show that

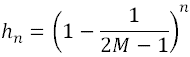

and

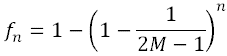

Comparing this result to the expression for the variance after *n* generations (Eq. 2), we notice that clonality within the cell is just a normalized mitochondrial variance between host cells,

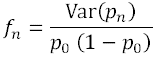

